# From silicone gel bleed chemistry to skeletal muscle and lipid alterations: clinical and *in vitro* evidence

**DOI:** 10.64898/2026.08.22.746421

**Authors:** N. Couturier, E. Randrianaridera, Cuong Le, H. Mutlu, I. Pluvy, F. Monnien, F. Bibeau, K. Anselme, A. Ponche, I. Brigaud

## Abstract

Musculoskeletal symptoms are frequently reported following silicone breast implantation. However, the biological mechanisms linking implant-derived silicone exposure to skeletal muscle alterations remain poorly understood, partly because the biological effects of silicone have long been debated in the context of its biocompatibility. Here, we chemically characterized the low-molecular-weight fraction of the breast implant silicone exposome, readily released from implant gel through “gel bleed”, and investigated its potential biological consequences using an integrated approach combining analytical chemistry, clinical transcriptomics and histology, and controlled *in vitro* muscle experiments. Transcriptomic analyses of periprosthetic tissues associated with silicone implant rupture revealed unexpected myogenic and neuromuscular signatures in tissue conventionally regarded as predominantly fibrous, together with alterations in lipid metabolism and transport. These findings were supported by histological evidence of close interactions between periprosthetic tissue and skeletal muscle. Chemical analysis of the implant-gel extract detected linear siloxane L2 and cyclic siloxanes D3–D8, with tentative assignment of D9. *In vitro*, C2C12 cells exposed to the implant-gel extract showed up to 30% reduced viability and decreased expression of key neuromyogenic genes. Together, these findings provide convergent chemical, clinical, and experimental evidence that low-molecular-weight constituents of the breast implant silicone exposome may constitute a biologically active exposure capable of affecting skeletal muscle. The associated alterations in lipid metabolism and transport further provide a mechanistic framework for investigating the cellular handling and potential tissue distribution of hydrophobic silicone-derived species. These findings position silicone gel bleed as a biologically relevant source of chemical exposure rather than solely a material-integrity phenomenon.

**Highlights:**

- Histological analysis revealed skeletal muscle areas closely integrated with periprosthetic fibrotic tissue.
- Patient-derived RNA sequencing reveals that silicone exposure exerts biological effects beyond inflammation, directly impacting both myogenic and lipid pathways.
- A gel implant-conditioned medium, chemically characterized by GPC and GC-MS, was used as an *in vitro* silicone exposure model and revealed a heterogeneous LMWS profile comprising L2 and cyclic siloxanes D3–D8, with D9 tentatively identified.
- The viability of differentiating muscle cells *in vitro* is altered by up to 30% by implant-derived LMWS.
- Silicone exposure induces muscle transcriptomic alterations *in vitro*, recapitulating key RNA-sequencing signatures observed in patient tissues, even in the absence of an inflammatory environment
- RNA-seq analysis uncovered dysregulation of lipid metabolism genes, including key lipoprotein markers, suggesting that silicone exposure may disrupt endogenous lipid transport pathways in periprosthetic tissues.

**Graphical Abstract:** 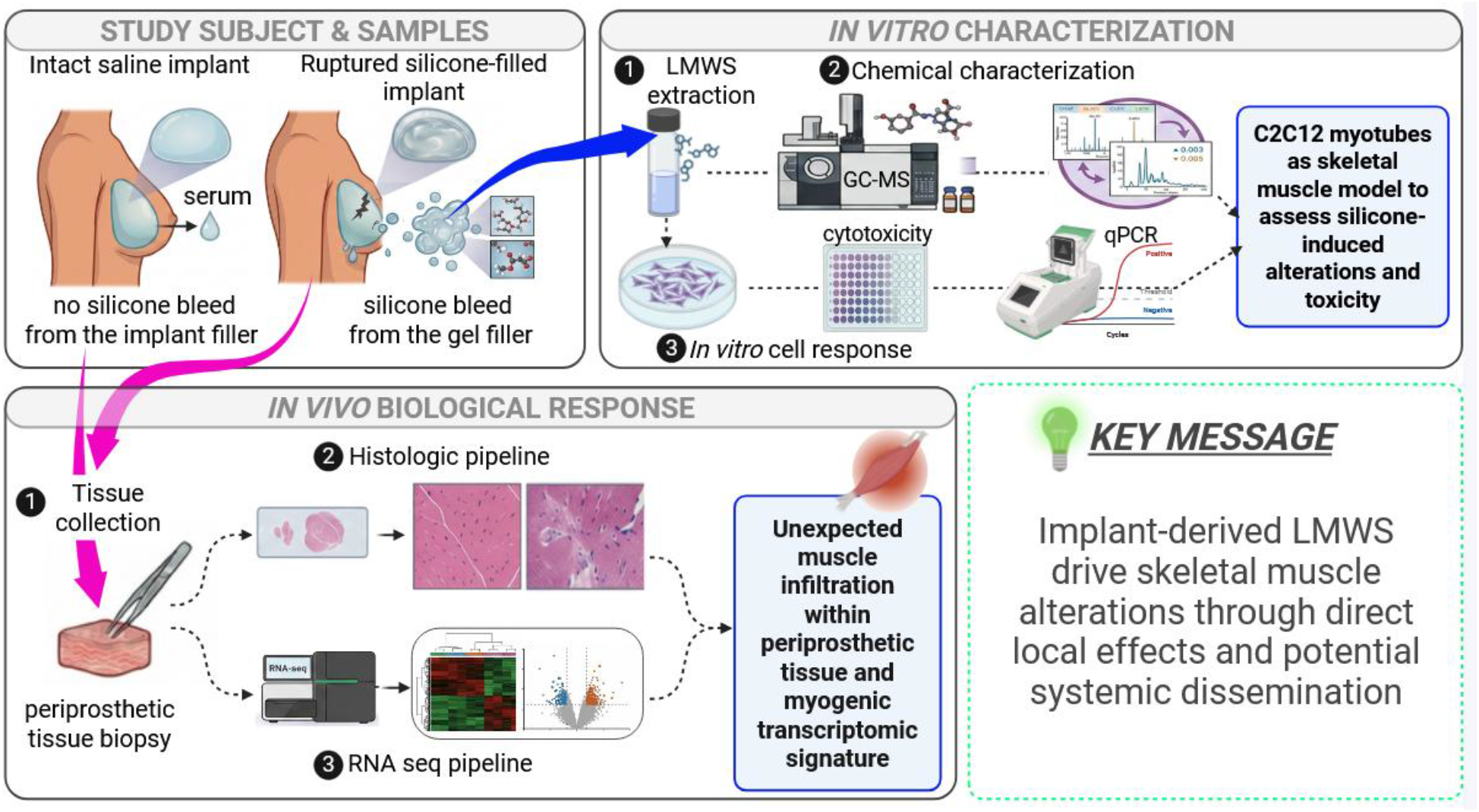

## Introduction

Globally, an estimated 35 million individuals have received breast implants, highlighting their extensive use in aesthetic and reconstructive surgery[1]. In 2024 alone, 1.66 million augmentation procedures were performed for cosmetic indications, establishing breast implantation as one of the most common aesthetic surgical procedures worldwide. That same year, approximately 342,000 breast implant removal interventions were conducted, representing a 65.3% increase compared with 2020. Although this increase may reflect growing clinical and public awareness of breast-implant-associated complications[2], the available procedural statistics do not establish the reasons for explantation. Among patients undergoing explantation because of implant-related systemic symptoms, approximately 82% have been reported to experience partial or complete symptom improvement following implant removal[3]. However, this estimate derives from observational studies and should be interpreted in the context of the studied patient populations and outcome definitions.

Although silicone has long been considered a biologically inert biomaterial, increasing evidence demonstrates that silicone implants participate in persistent host-material interactions involving immune activation, inflammation, and tissue remodeling[4]. Silicone exposure may occur through implant rupture, “gel bleed”, or degradation and mechanical wear of the implant shell, resulting in silicone accumulation in adjacent periprosthetic tissues[4] and, in some reports at distant anatomical sites[3,5,6]. Gel bleed is defined as the gradual permeation of low molecular weight siloxanes (**LMWS**) and related silicone-derived species from the implant gel across the elastomer shell. Accordingly, implant rupture represents a distinct failure mode that can produce substantially greater and less controlled gel release and is considered the extreme case of gel permeation[7].

*In vivo,* evidence of silicone release primarily derives from histological analyses of periprosthetic tissues, which consistently reveal silicone droplets, often located within macrophage cytoplasm, indicating phagocytosis of leaked material[4,7]. Recent studies have reported up to one million silicone particles per periprosthetic tissue sample, even in the latest designs, highlighting that silicone gel bleed remains a contemporary and clinically relevant phenomenon[8]. Furthermore, LMWS have been detected beyond the periprosthetic tissue to mammary lymph nodes and, in some cases, to visceral organs such as the liver, spleen, or brain[5,7]. Although these observations suggest that silicone-derived compounds may disseminate beyond the implantation site, the mechanisms, extent, and clinical significance of this distribution remain incompletely understood. Yet, whether persistent inflammation is directly related to systemic LMWS dissemination or reflects multiple interacting biological processes remains unresolved[3,7,9]. While immune complications from local silicone exposure are well documented[7,10–12], the physicochemical mechanisms driving LMWS migration and their potential contribution to distant tissue alterations remain poorly documented.

Since their introduction in the 1960s, silicone breast implants have been linked to systemic disease, with over 100 symptoms reported in the literature and collectively termed breast implant illness (**BII**). BII is a heterogeneous syndrome encompassing fatigue, myalgia, arthralgia, cognitive dysfunction, and sicca symptoms. Remarkably, with a reported prevalence of 28% in implanted patients, musculoskeletal symptoms, including pain, weakness, stiffness, cramps and debilitating fatigue, dominate the clinical presentation of BII[3]. However, the reported prevalence varies considerably between studies owing to differences in patient populations, study design, and diagnostic criteria. These symptoms can severely impair quality of life and, in some case, lead to physical disability. Although BII is not currently recognized as a distinct medical diagnosis, its clinical presentation substantially overlaps with autoimmune/inflammatory syndrome induced by adjuvants (**ASIA**), in which silicone has been proposed to act as an immunological adjuvant, triggering sustained immune response and paving the way to the development of breast implant associated diseases[13,14]. Despite being suspected for decades, the pathophysiological mechanisms linking silicone exposure to systemic manifestations remain unclear. Three major barriers hinder progress in understanding BII and silicone-induced systemic effects: (1) the absence of clinically relevant experimental models that fully reproduce long-term human implantation (2) the delayed and highly variable onset of symptoms, which complicates the identification of initiating events and (3) human tissue access is limited to the periprosthetic tissue, thereby preventing longitudinal characterization of disease progression and comprehensive analysis of distant organs, including skeletal muscle. Consequently, despite long-standing interest in silicone implant safety, the molecular pathways linking chronic silicone exposure, immune dysregulation, and musculoskeletal symptoms remain unidentified. In a nutshell, despite long-standing interest in silicone implant safety, the mechanisms underlying breast-implant related systemic myotoxic effects remain incompletely understood and the clinical entity itself remains controversial[3,15,16].

Our previous work established persistent immune activation, inflammation, and tissue remodeling associated with silicone exposure, including in cases of clinically silent implant rupture[4]. Building on these findings, we sought to investigate whether silicone exposure may also be associated with biological pathways beyond immune-related responses. We focused on two unexpected molecular signatures: (i) a myogenic transcriptional profile, offering new insights into potential mechanisms underlying silicone-associated muscle alteration, and (ii) dysregulation of lipid metabolism pathways, suggesting a possible impact of silicone exposure on endogenous lipid transport systems. To further investigate the potential biological relevance of the myogenic signature, we developed and chemically characterized an *in vitro* model based on the low-molecular-weight fraction released from silicone breast implant gel through “gel bleed”. Using commercially available breast implants, we chemically characterized the resulting implant-derived low-molecular-weight silicone (**LMWS**) profile and exposed differentiating C2C12 myoblasts to the gel-bleed extract to assess cell viability and the expression of selected myogenic and neuromuscular genes. Rather than seeking to establish causality in patients, this complementary *in vitro* model was designed to determine whether implant-derived LMWS can trigger biological responses in skeletal muscle cells under controlled experimental conditions.

Collectively, this work provides experimental evidence, across clinical tissues and an *in vitro* model, that the low-molecular-weight fraction of the breast implant silicone exposome, readily released from implant gel through gel bleed” represents a biologically relevant exposure rather than merely a material-integrity phenomenon. In this study, we demonstrate that silicone exposure is associated with myogenic dysregulation and altered lipid homeostasis in periprosthetic tissues following implant rupture, and provide *in vitro* evidence that exposure of muscle cells to the implant-derived fraction alters selected myogenic and neuromuscular genes identified in the clinical analysis, even in the absence of exogenous inflammatory stimulation. Together, these findings extend the biological scope of silicone exposure beyond the conventional scope of immune-related responses and provide a new materials-to-biology framework for investigating the interactions of low-molecular-weight silicone species with skeletal muscle and endogenous lipid-handling pathways.

## Methods

### RNA sequencing

#### (i) Tissue sampling and data collections

Periprosthetic tissues were obtained from patients previously enrolled in our institutional biobank protocol[4], and managed by the Tissue Bank of Franche-Comté (BB-0033-00024, University Hospital of Besançon, France). The study protocol was reviewed and approved by the localScientific Board and the Institutional Review Board of the Tumorothèque Régionale de Franche-Comté, the regional biobank of the University Hospital of Besançon (registration no. BB-0033-00024; project approval no. F967-INFLAMA; collection declaration no. DC-2014-2086; IRB: NCT06414785), and all participants provided written informed consent. Histological sections were collected whenever available. Tissue samples intended for RNA sequencing were immersed in RNAlater immediately after explantation and stored at −20 °C until processing. After anonymization, demographic, clinical, and implant-related data were retrieved, including patient age, implant manufacturer, surface topography, implantation duration, indication for explantation, and implant integrity (detailed in[4]). Implant integrity was determined intraoperatively by macroscopic examination of the explanted device. The present study was restricted to ruptured silicone gel-filled breast implants compared to saline-filled intact implants. To minimize potential confounding by pre-existing inflammation, only non-malignant Baker grade I capsules from otherwise healthy patients were included.

#### (ii) RNA-Sequencing

This study reanalyzes the transcriptomic dataset previously described in the context of inflammatory processes[4]. Briefly, total RNA was extracted from human periprosthetic capsular tissues adjacent to intact saline-filled implants (n = 3) or ruptured silicone gel-filled implants (n = 8), using the RNeasy Fibrous Tissue Mini Kit (Qiagen), following mechanical homogenization (Precellys Evolution, Bertin Instruments). Stranded mRNA libraries were prepared at the GenomEast platform (IGBMC, France) from 100 ng of total RNA using the Illumina Stranded mRNA Prep Ligation kit and IDT for Illumina RNA UD Indexes. Libraries were sequenced on an Illumina NextSeq 2000, and raw data were processed with Cutadapt (v4.2) before alignment to the GRCh38 human genome (Ensembl release 110).

#### (iii) Differential gene expression analysis

Differential gene expression analysis was conducted using the DESeq2 package (v1.34.0) in R, building upon the original primary analysis from[4]. Differential gene expression was evaluated by comparing periprosthetic tissues exposed to extensive silicone gel bleed following implant rupture with periprosthetic tissues surrounding intact saline-filled implants, which served as a gel bleed-free control. Differentially expressed genes (**DEGs**) were selected based on a combination of significance threshold of p-value < 0.05 and an absolute fold change (|log2FC| > 1). The distribution and magnitude of differential expression were visualized using Volcano plots generated with OriginLab 2024. Upregulated and downregulated DEGs were summarized using bar plots to illustrate the overall transcriptional changes. The complete DEG dataset was subsequently used for *functional enrichment analyses*. Gene Ontology (**GO**) enrichment analyses were performed across the three main GO domains: Biological Process (**BP**), Cellular Component (**CC**), and Molecular Function (**MF**). In addition, Kyoto Encyclopedia of Genes and Genomes (**KEGG**) pathway enrichment analysis was performed to identify significantly affected biological pathways. GO and KEGG pathway enrichment analyses were conducted using g:Profiler (e114_eg62_p19_27110d83)[17]. Significant terms were identified with a p-value threshold of 0.05 and corrected for multiple testing using the Benjamini-Hochberg FDR. The background was set to all human genes (ENTREZGENE_ACC).

#### (iv) Functional categorization of enriched GO Biological Process terms

To facilitate biological interpretation of enriched GO biological process (BP) terms, a keyword-based semantic classification was applied to systematically group significant terms into broader functional categories. Namely, GO:BP terms were categorized into four major functional groups based on predefined keyword sets and expert-driven biological relevance: (1) **Inflammation**: All terms explicitly containing *“inflammatory”*, *“immune”*, *“defense”*, *“cytokine”*, *“leukocyte”*, *“lymphocyte”*, *“macrophage”*, *“neutrophil”*, *“T cell”*, *“B cell”*, *“natural killer cell”*, *“antigen”*, or *“wound healing”*, including regulatory processes (e.g., *“regulation of immune response”*) and responses to biotic/abiotic stimuli (e.g., *“response to bacterium”*). (2) **Skeletal Muscle Processes**: terms containing *“muscle”*, *“sarcomere”*, *“myo-”*, *“actin”*, *“striated”*, *“myofibril”*, or *“contraction”.* (3) **Neuronal Signaling**: terms containing *“signal transduction”*, *“synaps”*, *“neuro-”*, *“receptor”*, *“action potential”*, *“membrane depolarization”*, or *“calcium-mediated signaling”*. (4) **Lipid Metabolism**: terms containing *“lipid”*, *“fatty acid”*, *“sterol”*, *“cholesterol”*, *“phospholipid”*, *“sphingolipid”*, or *“glycerolipid”*.

#### (v) Focused analysis of key functional categories

Given their biological relevance to silicone-associated pathophysiology, Skeletal Muscle Processes and Lipid Metabolism were selected for in-depth analysis. These broad categories were further subdivided into specific subcategories to enable a higher-resolution interpretation of the enriched processes. Finally, bubble plots, integrating enrichment significance and biological representation were used to visualize the contribution of specific functional themes within the transcriptomic dataset. Bubble plots were generated online using SRplot package[18].

### Histology and image acquisition

Complete methodological details and sampling are provided in[4]. Briefly, we obtained serial 4 μm-thick sections from formalin-fixed, paraffin-embedded breast capsule samples using a microtome and stained them with hematoxylin and eosin (H&E). Microscopic observations were performed using a Leica DM2000 light microscope equipped with 20×, 40×, and 63× objectives. Micrographs were captured with a digital camera (LAS Version 4.2) and assembled using Adobe Photoshop CC 2019. Histological analysis was conducted by a certified histologist at Novotec Institute (Bron, France) under blinded conditions.

### *In vitro* model of silicone gel bleed

#### (i) Commercial silicone breast implant gel source

The silicone gel was extracted from a new, unimplanted SEBBIN breast implant (ref. LSC 55415; Naturgel™), featuring a smooth (non-textured) surface, round shape, and 415 mL of highly cohesive, medical-grade silicone gel encapsulated in a flexible polydimethylsiloxane (PDMS) elastomer shell. All manipulations were performed under UV-free conditions to prevent cross-linking alterations. Manufacturer specifications are available in the SEBBIN instructions for use (ISO 14607 compliant; www.sebbin.com/ifu/).

#### (ii) LMWS extraction protocol

To simulate *in vivo* gel bleed, we adapted a protocol inspired by Dijkman et al[19]. We aseptically incised with a sterile scalpel under a laminar flow hood and collected 5 g gel aliquots. Samples were stored dry at stable room temperature in the dark until extraction. LMWS extraction protocol was adapted from adapted from Dijkman (2023)[19]. For extraction, 5 g gel aliquots were statically incubated in 20 mL C2C12 serum-free cell culture medium (DMEM; Gibco, Cat. No. 41966029) at 37°C for 10 days. The resulting LMWS-containing solutions were either directly use for downstream chemical characterization (GPC, GC-MS) or applied to cellular assays. All procedures were performed under sterile conditions, and endotoxin levels in LMWS solutions were verified to be < 0.1 EU/mL using a limulus amebocyte lysate (LAL) assay (ToxinSensor™ Gel Clot Endotoxin Assay Kit (cat. no. L00351, GenScript, Piscataway, NJ, USA).

#### (iii) LMWS characterization: Gel Permeation Chromatography (GPC)

Siloxane species were extracted from gel-infused medium via liquid-liquid extraction using dichloromethane (DCM stabilized with ethanol, Carlo Erba, >99.95 %) at a 1:1 volume ratio (10 mL medium:10 mL DCM). After solvent evaporation, the organic residue was reconstituted in 1.0 mL tetrahydrofuran (THF, stabilized with butylated hydroxytoluene, >99.8 %) and analyzed by GPC using hexamethylcyclotrisiloxane (D3, ABCR GmbH, > 95 %), octamethylcyclotetrasiloxane (D4, Wacker, >99.9 %), and dodecamethylcyclohexasiloxane (D6, Supelco analytical standard, >97 %) as reference standards. Molecular weight analysis was performed using size exclusion chromatography (SEC) on an Agilent 1260 Infinity system equipped with an autosampler, a column set comprising a guard column (50 × 7.5 mm) and two analytical ResiPore columns (300 × 7.5 mm; particle size: 3 μm; porosity: 2 μm; Polymer Laboratories), a G1314B variable wavelength detector (280 nm), and a G7800A multidetector suite (refractive index + viscosimeter). Measurements were conducted at a flow rate of 1 mL/min at 35 °C using THF as the eluent. Calibration employed linear polystyrene standards (EasiVial, Agilent; range: 162 – 3.64 × 10⁵ g/mol). Polymer samples were dissolved in THF at 2 mg/mL, filtered through a 0.2 μm filter, and analyzed using Agilent GPC/SEC software.

#### (iv) Gas Chromatography–Mass Spectrometry (GC-MS)

Extracts were analyzed using a Shimadzu QP2010 gas chromatograph coupled to a mass spectrometer. Chromatographic separation was performed on a SGE Analytical Sciences capillary column packed with BPX-5 stationary phase (25 m × 0.25 mm, 0.25 μm film thickness). The carrier gas was helium, used at the constant linear velocity of 25cm/s (corresponding to a pressure of 20.5 psi and at the column flow of 0.40 mL/min with an oven temperature of 40°C for our system). The injection volume was 1 µL in splitless mode, with an injector temperature of 250 °C. The transfer line temperature is maintained at 300 °C and the EI source temperature at 200 °C. The oven temperature program was as follows: an initial temperature of 40 °C held for 4 min, ramped at 7 °C/min to 150 °C, then ramped at 14 °C/min. to 260 °C, and held for 4.43min. The total analysis time is 32 min. MS detection was performed by Electronic Impact (EI) in full scan mode (m/z from 46 to 400) for identification, and in Selected Ion Monitoring (SIM) mode for quantification, targeting characteristic ions of siloxanes: hexamethyldisiloxane L2 (147 and 73), hexamethlycyclotrisiloxane D3 (207, 191, 133 and 96), octamethylcyclotetrasiloxane D4 (281, 193, 133 and 73) and cyclopentasiloxane D5 (355, 267 and 73). Dodecamethylcyclohexasiloxane (D6), tetradecamethylcycloheptasiloxane (D7), hexadecamethylcyclooctasiloxane (D8), octadecamethylcyclononasiloxane (D9) were identified by comparing mass spectra obtained in full scan mode with those of the NIST 2017 library; a minimum match factor of 89% was required for compound identification. Concentrations were estimated at the order-of-magnitude level by single-point external standardization using certified pure compounds (L2, D4, D5, D6; Sigma-Aldrich, ≥98%) at 50 ng/mL, corresponding to the trace-level quantification threshold of the analytical SIM method.

### *In vitro* assays: C2C12 cells as a myogenic biosensor to silicone exposure

#### (i) C2C12 cell culture and silicone exposure

##### Cell maintenance

C2C12 murine myoblasts were maintained in growth medium consisting of Dulbecco’s Modified Eagle Medium (DMEM; Gibco, Cat. No. 41966029) supplemented with 15% fetal bovine serum (FBS, Gibco, Cat. No. A5209501) and 1% penicillin–streptomycin (Sigma, Cat. No. P0781). Cells were passaged every other day using Trypsin-EDTA (Gibco, Cat. No. 25200-056) and maintained below 60% confluence to preserve myogenic potential. **For exposure experiments,** cells were seeded at 70,000 cells/cm² on fibronectin-coated 6-well plates (Corning Costar, Cat. No. 3516; Fibronectin coating concentration: 3 µg/mL, Sigma, Cat. No. F4759). After 24h, differentiation was induced by replacing growth medium with differentiation medium consisting of DMEM (Gibco, Cat. No. 41966029) supplemented with 2% horse serum (Gibco, Cat. No. 16050-130) and 1% penicillin–streptomycin. Control cells received differentiation medium prepared with native DMEM, while silicone-exposed cells were cultured in differentiation medium prepared with silicone-infused DMEM (see Section *in vitro model of silicone bleed*). Differentiation medium was replaced every other day, and cells were allowed to differentiate for 6 days.

#### (ii) Video microscopy

Time-lapse imaging was performed using an Incucyte SX1 live-cell analysis system (Sartorius) with a 20× objective, acquiring phase-contrast images every 2 hours. Images were acquired at a resolution of 0.62 µm/pixel (1408 × 1040 pixels).

#### (iii) RNA isolation and real-time quantitative PCR

Total RNA was isolated from C2C12 myotubes using a column-based extraction method (RNeasy Plus Micro Kit, Qiagen, Cat. No. 74034) according to the manufacturer’s instructions. For RT-qPCR, 500 ng of total RNA was reverse transcribed using the iScript Reverse Transcription Supermix for RT-qPCR (Bio-Rad, Cat. No. 1708841), and qPCR was performed using iTaq Universal SYBR Green Supermix (Bio-Rad, Cat. No. 1725124) on a CFX Connect Real-Time PCR Detection System (Bio-Rad), following the manufacturer’s protocol. All reactions were performed in duplicate. *Rpl0* and *Pol2* were used as housekeeping genes for normalization of gene expression. Primer sequences are listed below, **Tab.1**. Experiments were performed independently in triplicates (n=3).

**Table. 1.**
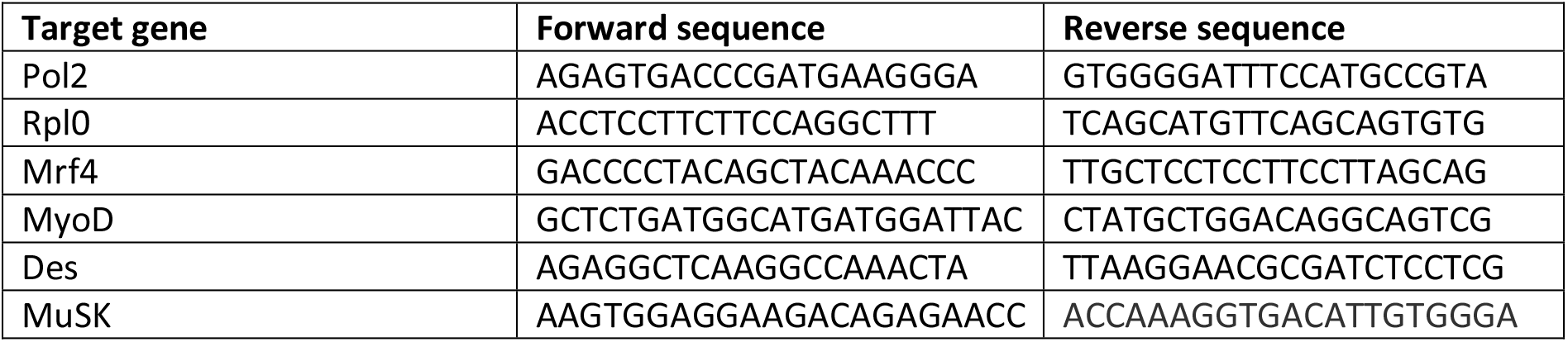
qPCR primer sequences.

#### (iv) Cell viability assay

After 6 days of differentiation, cell viability was assessed using the Alamar Blue Assay. C2C12 myotubes were incubated for 1 h at 37 °C and 5% CO₂ in differentiation medium supplemented with 10% Alamar Blue reagent (Invitrogen, Cat. No. DAL1100). Fluorescence was measured using a SpectraMax iD3 microplate reader (Molecular Devices) at excitation/emission wavelengths of 545/590 nm.

#### (v) Statistical analysis

Statistical analyses were performed using R software. Comparisons between control and permeates-treated samples were performed using two-tailed paired Student’s t-tests on data from three independent biological experiments (n = 3). For RT-qPCR analyses, statistical tests were performed on ΔCt values, whereas relative gene expression was calculated using the 2^−ΔΔCt^ method for graphical representation, with the control condition normalized to 1. Data are presented as individual biological replicates. Statistical significance was defined as *p* < 0.05 and indicated as follows: *p* < 0.05 (\**), p < 0.01 (**), p < 0.001 (\*\*\**), and *p* < 0.0001 (****).

## Results

### 1. Characterization of gel bleed-derived LMWS from commercial breast implant

To identify silicone-derived compounds that are readily released into an aqueous environment under the applied experimental conditions, we established an *in vitro* extraction model using commercially available silicone breast implant gel. The resulting siloxane-derived compound blend released into the aqueous medium was characterized by GPC, **Fig. 1A**. Comparison of chromatograms from gel-infused medium with those of authenticated standards revealed a peak consistent with D3 eluting at approximately 20.5 min (**Fig. 1A**). A less-resolved signal was also detected at the D4 retention time (t_R_ = 20.1 min), although partial overlap with matrix components prevented unambiguous interpretation. No distinct peak was observed at the expected retention time of D6, suggesting either limited recovery under these analytical conditions or interaction with the stationary phase. However, a broad signal eluting before the D6 window pointed to the presence of higher-molecular-weight siloxane species, consistent with the presence of silicone-derived species with larger apparent hydrodynamic volumes (**Fig. 1A**). Whether the GPC analysis indicated the presence of LMWS derived species in the aqueous extract, it did not allow unequivocal structural assignment. Given the limitations of GPC for compound identification and its primarily relative molecular-weight information, the extracts were further analyzed by GC-MS for structural identification and semi-quantitative characterization (**Fig. 1B**).

**Figure 1.**
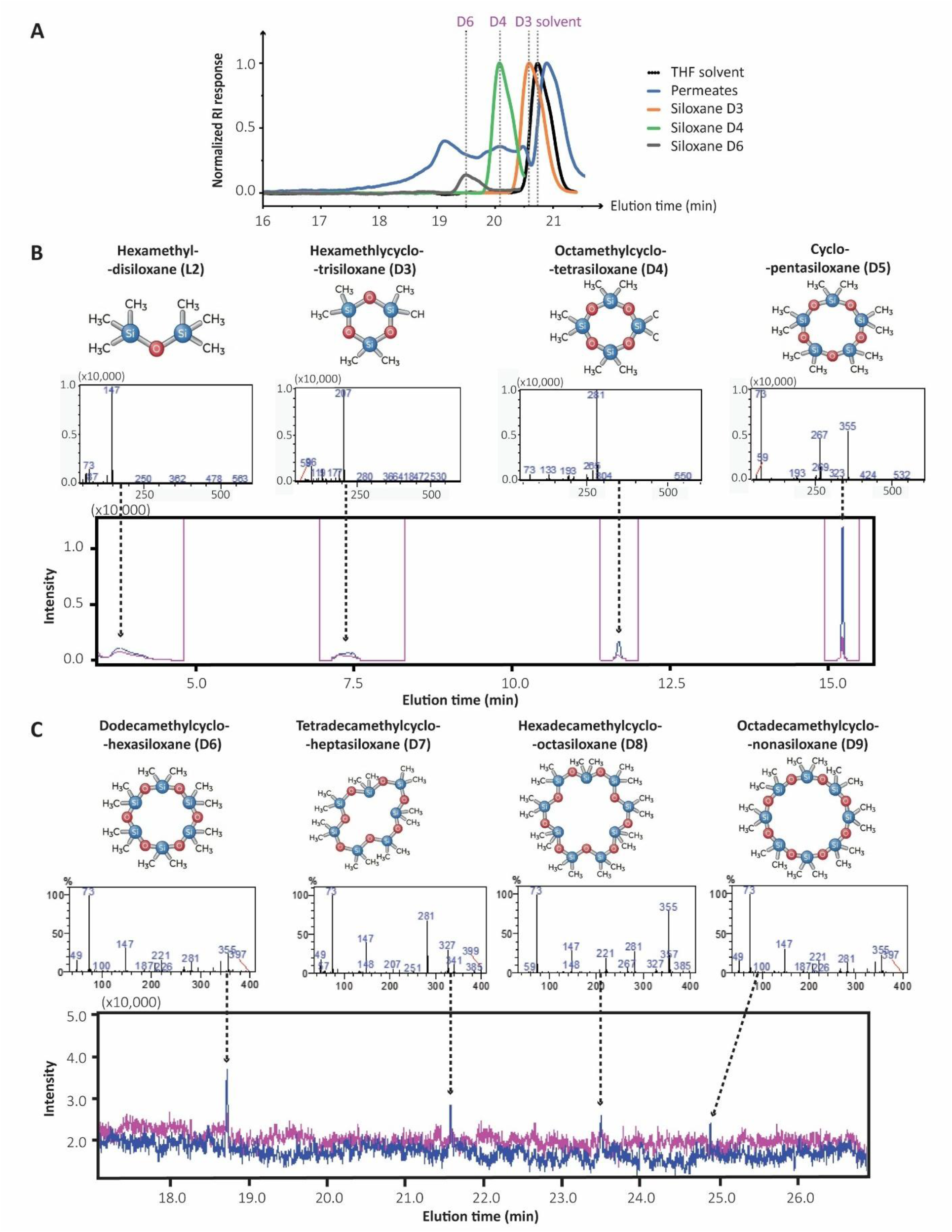
Identification and characterization of LMWS in silicone gel bleed extracts. (**A**) GPC chromatograms of gel-infused medium (blue), THF solvent (black), and siloxane standards: D3 (orange), D4 (green), D6 (purple). Peaks at ∼17.5 min (D6), ∼18.5 min (D4), and ∼20.5 min (D3) confirm the presence of these cyclic siloxanes in permeates. (**B**) GC-MS analysis of smaller cyclic siloxanes in gel infused (blue) or control (pink) medium: chemical structures (top), mass spectra with characteristic m/z fragments (middle), and chromatograms (bottom) for L2, D3, D4, and D5. Dotted lines indicate peak assignments. (**C**) GC-MS analysis of larger cyclic siloxanes in gel infused (blue) or control (pink) medium: chemical structures (top), mass spectra (middle), and chromatograms (bottom) for D6, D7, D8, and D9. Later elution times reflect uncharacterized higher molecular weights.

Comparison with identically processed control medium (without gel) confirmed active LMWS release from silicone gel, marked by elevated peak intensities for L2, D3, D4, and D5 (A_infused medium_ vs A_control medium_ = 5377 vs 2220, 4512 vs 934, and 5388 vs 881 a.u. respectively, **Fig. 1B**), consistent with release of these compounds from the silicone gel. Low control signals align with reports of volatile cyclosiloxanes as common laboratory contaminants originating from solvents, laboratory consumables and personal care products[20–22], enabling distinction of implant-derived compounds from background. Full-scan GC-MS chromatograms revealed additional late-eluting compounds (**Fig. 1C**). Mass spectral comparison with the NIST 2017 spectral library identified these as D6, D7 and D8 (A_infused medium_ vs A_control medium_ = 5956 vs 740, 2479 vs 468, and 2014 vs 881 a.u. respectively). An additional chromatographic peak eluting at 24.9 min displayed a fragmentation pattern consistent with a higher molecular weight cyclosiloxane. Although the *m/z* < 400 acquisition range precluded definitive identification its retention time and fragmentation profile were consistent with D9 which was therefore considered a tentative assignment. The GC-MS profile extends beyond the volatile cyclosiloxanes typically studied in breast implant research (D3-D5), also detecting the lighter L2 together with cyclic siloxanes D3–D8 and a putative D9 under the experimental conditions employed (**Fig.1**).

### 2. Transcriptomic signatures in silicone-exposed periprosthetic tissues

#### 1.1 Global transcriptomic dysregulation in silicone-exposed periprosthetic tissues extends beyond inflammation

Differential gene expression analysis between periprosthetic tissues from ruptured silicone gel-filled implants and intact saline-filled controls (serving as a gel bleed-free reference), revealed broad transcriptional dysregulation, with 3,993 differentially expressed genes (**DEGs**), comprising 1,629 upregulated and 2,364 downregulated genes (**Fig. 2A-B**). To gain insight into the biological processes associated with these DEGs, we conducted Gene Ontology (GO) enrichment analysis across Biological Process (BP), Cellular Component (CC), and Molecular Function (MF), as well as KEGG pathway enrichment (**Fig. 2C**). The 1,971 significant GO:BP terms identified were then categorized into broad functional groups based on keyword-based semantic classification. Strikingly, inflammation-related terms accounted for only 12.5% of the total, while neuronal signaling (9.5%), muscle-related processes (9%), and lipid metabolism (6.5%) collectively represented 25% of the transcriptomic response. With, non-inflammatory processes outweighing inflammatory ones, our results underscore the complexity of silicone-induced biological mechanisms, which thus extend far beyond immune activation solely. Given that the inflammatory component of this response has been previously described[4], we focused our analysis on neuromuscular and lipid metabolism pathways, two categories of particular biological relevance to silicone-associated pathophysiology and associated silicone systemic transport.

**Figure 2.**
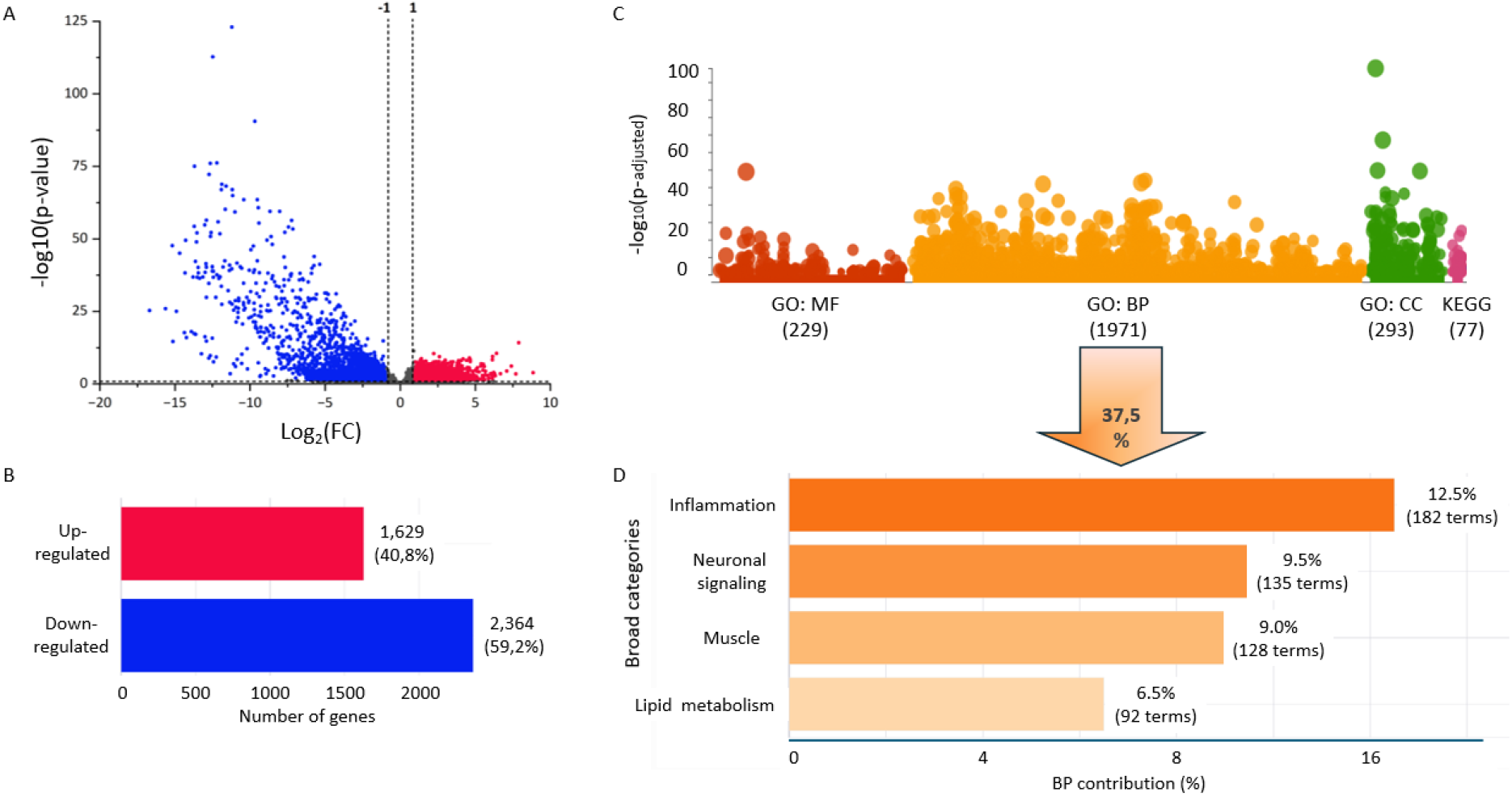
Effect of silicone exposure following breast implant rupture on gene expression in periprosthetic tissue. (**A**) Volcano plot and (**B**) Bar chart showing the distribution of up- and down-regulated genes, (**C**), Complete GO enrichment analysis profile with gene sets colored into categories: GO MF (Molecular Function, red), GO BP (Biological Process, orange), GO CC (Cellular Component, green) and KEGG (pink), (**D**), Proportional contributions of biological programs (% by category-associated term numbering), focusing on inflammation, skeletal muscle processes, neuronal signaling, and lipid metabolism.

#### 2.2 Unexpected integration of skeletal muscle structures and myogenic transcriptional signatures in periprosthetic tissues

Periprosthetic tissue is conventionally described as predominantly fibrous[23], making the identification of a prominent myogenic and neuromuscular transcriptomic signature unexpected. We therefore examined the histological composition of the periprosthetic capsules to determine whether skeletal muscle was physically represented within the tissue analyzed by RNA sequencing. Histological examination of the periprosthetic capsules revealed recurrent presence of skeletal muscle bundles intimately integrated within the fibrotic periprosthetic tissue (**Fig. 3A**). Rather than being anatomically separated from the fibrotic periprosthetic tissue, skeletal muscle fibres were found embedded within the capsule indicating a close and entangled spatial organization between the periprosthetic tissue and adjacent skeletal muscle. Higher magnification further confirmed the close association between muscle fibres and connective tissue throughout the periprosthetic tissue (**Fig. 3Ad**). The presence of skeletal muscle components within silicone-exposed periprosthetic capsules following implant rupture provided an unusual opportunity to investigate associated transcriptional alterations in muscle tissue. We therefore analyzed our RNA-seq dataset with a focus for genes and pathways related to skeletal muscle biology. Differential expression analysis identified a substantial subset of differentially expressed genes (DEGs) associated with muscle-related biological processes. Muscle-related genes accounted for 503, exclusively downregulated, DEGs (**Fig. 3B**), revealing a strongly impacted transcriptional program. Functional classification showed that genes involved in muscle development and morphogenesis represented the largest category (28%), followed by regulation of muscle processes (22%) and muscle contraction and function (18%) (**Fig. 3C**). Consistent with these observations, Gene Ontology enrichment analysis identified biological processes associated with skeletal muscle development, differentiation, satellite cell activation, muscle hypertrophy and myofibril assembly (**Fig. 3D**). Interestingly, several canonical regulators of myogenic commitment and muscle differentiation, and multiple myosin heavy-chain isoforms, were among the dysregulated genes (**Fig. 3B**; gene list detailed in **supplementary table S1**), evidencing a profound remodelling of the myogenic program following silicone exposure.

**Figure 3.**
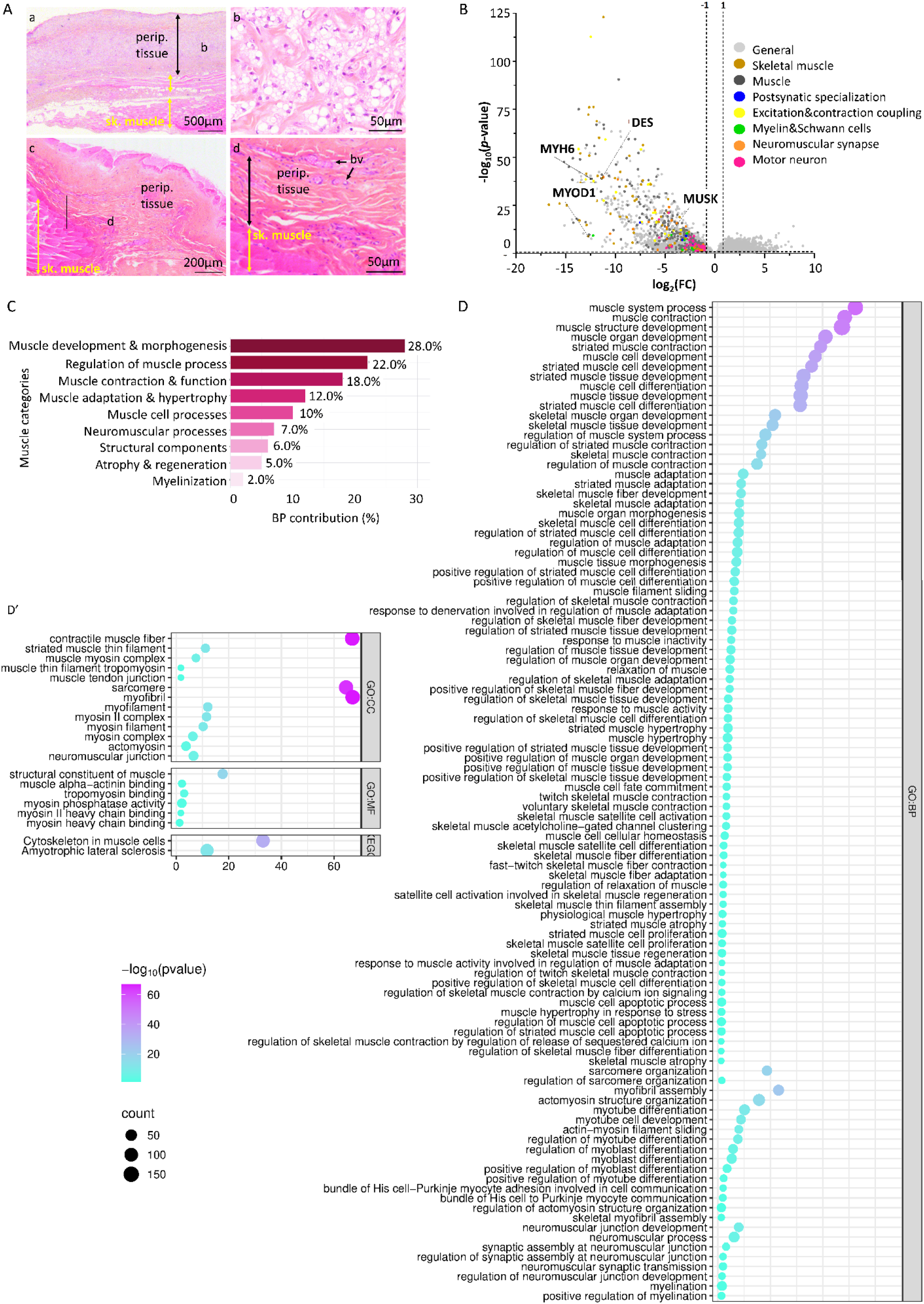
Silicone exposure following breast implant rupture alters muscle-related gene expression in periprosthetic tissues. (**A**) Representative hematoxylin and eosin (H&E)-stained periprosthetic tissues showing close integration of skeletal muscle bundles within the fibrotic capsule, irrespective of implant membrane integrity (**Aa–Ab**, ruptured implants; **Ac–Ad**, intact implants). (**Ab**) Palisading granuloma composed of foamy histiocytes associated with silicone exposure following implant rupture, consistent with a foreign body reaction to silicone particles. (**Ad**) Representative periprosthetic tissue from an intact implant showing normal tissue architecture with blood vessels and skeletal muscle integration. Perip. tissue, periprosthetic tissue; bv, blood vessel; Sk. muscle, skeletal muscle. (**B**) Volcano plot showing differentially expressed muscle-related genes, color-coded according to functional categories. (**C**) Relative contribution of muscle-related functional categories, expressed as the percentage of differentially expressed genes assigned to each category. (**D-D’**) Bubble plot of significantly enriched muscle-related biological pathways. Bubble color represents statistical significance (−log₁₀ P value), and bubble size corresponds to the number of associated genes.

**Supplemental data S1:**
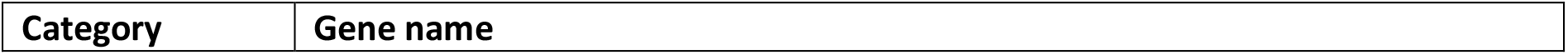
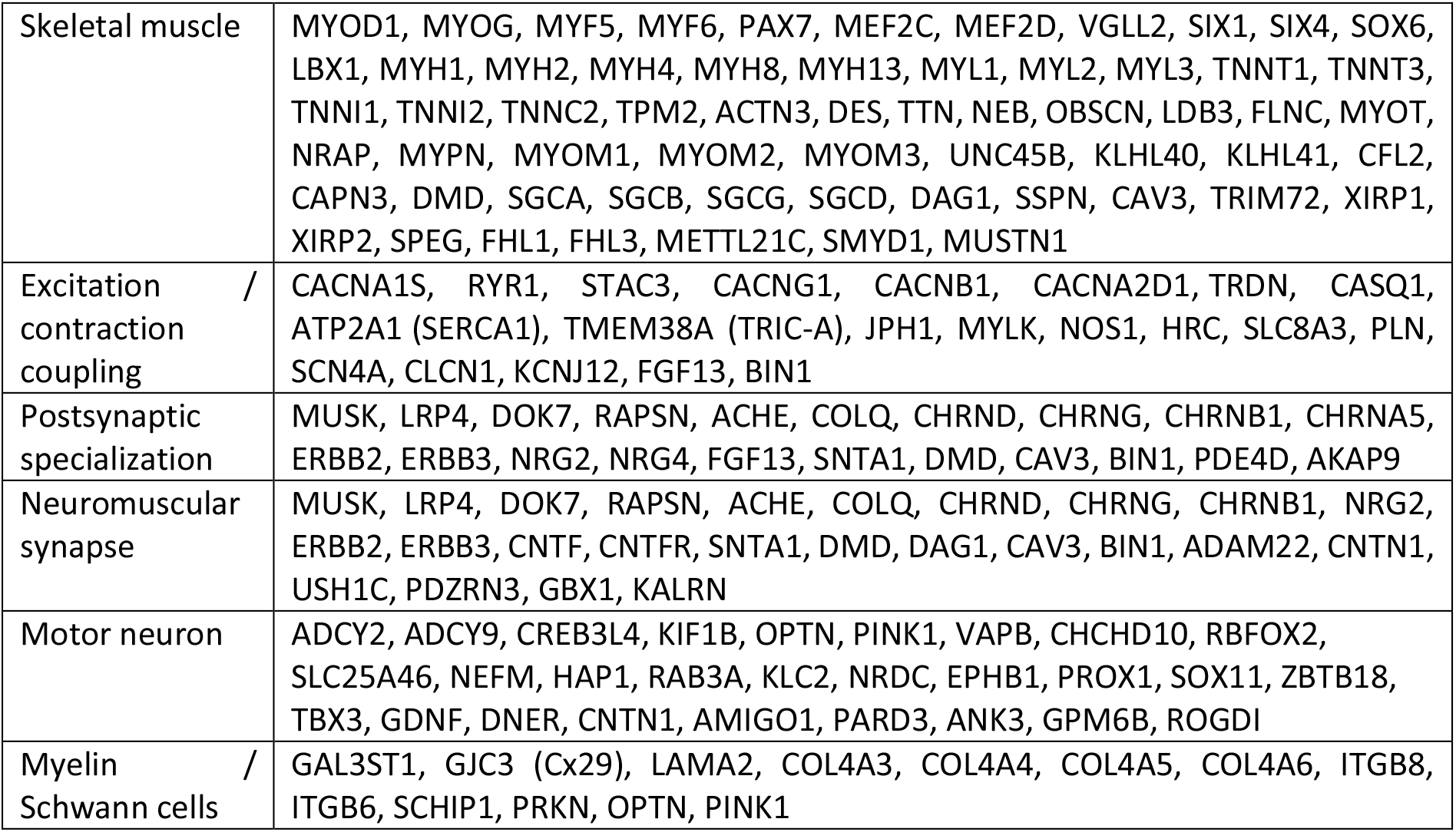
Gene list related to neuromuscular functions.

Beyond developmental pathways, the transcriptomic signature extended to genes controlling muscle contractile activity. Numerous dysregulated genes encoded essential components of the excitation-contraction coupling machinery, including calcium handling proteins (*RYR1*, *CACNA1S*, *ATP2A1*, *CASQ1*), ion channels (*SCN4A*) and sarcomeric structural proteins, while GO enrichment highlighted skeletal muscle contraction, contractile muscle fibre and sarcomere organization among the most significantly enriched terms (**Fig. 3D**). At the cellular component level, enrichment of myofibril, sarcomere and myosin complex further supported alterations affecting the contractile apparatus.

Remarkably, the molecular signature also extended beyond muscle fibres themselves. Dysregulated genes involved in neuromuscular junction organization and synaptic specialization, including *MUSK* and *LRP4* and additional neuromuscular junction-associated genes, were identified together with enriched biological processes related to neuromuscular junction development, synapse assembly and synaptic transmission (**Fig. 3A and 3D**). In parallel, genes associated with Schwann cells and myelination were also differentially expressed, suggesting that pathways involved in peripheral nerve support and nerve-muscle communication are represented within the periprosthetic transcriptome. This integrated neuromuscular signature was further supported by KEGG enrichment of the Amyotrophic Lateral Sclerosis pathological pathway, which includes numerous genes involved in neuromuscular junction integrity, axonal communication and muscle homeostasis. Collectively, these findings demonstrate that the transcriptional alterations observed in periprosthetic tissues encompass multiple levels of skeletal muscle biology, ranging from myogenic differentiation and contractile function to neuromuscular organization.

### Silicone permeates induce myotoxic effects and impair the myogenic program in differentiating C2C12 myotubes

To investigate whether silicone-derived compounds released from the gel affect skeletal muscle cells, C2C12 myoblasts were differentiated for 6 days in the absence (control) or presence of implant-derived silicone permeates, followed by assessment of metabolic activity and the expression of key myogenic markers (**Fig. 4**). Silicone permeate exposure markedly reduced cellular metabolic activity, as determined by the Alamar Blue reduction assay. Compared with control cells, exposed cultures exhibited a significant decrease in Alamar Blue signal from 62.78 ± 4.29% to 43.80 ± 3.13% (Fig. 4B), corresponding to an average 30.2% reduction (**Fig. 4B**). These findings indicate that silicone permeates significantly impair myotubes’ viability *in vitro*. Time-lapse video microscopy further confirmed the cytotoxic effect (**supplemental movies S2-S3**). Interestingly, although silicone permeates did not prevent initial myoblast fusion and myotube formation, permeate-exposed myotubes progressively underwent cell death during the differentiation period, whereas control cultures remained stable (**Fig. 4A-A’** and **supplemental movies S2-S3**). Then, to assess whether silicone exposure perturbed myogenic differentiation, we evaluated the expression of key myogenic regulatory genes highlighted by our RNA-seq analysis (**Fig. 3A**). Silicone permeates exposure markedly reduced the expression of muscle-associated markers including **MyoD1** (0.78 ± 0.06-fold; **Fig. 4C**), **Myh6** (0.11 ± 0.03-fold relative to control levels; **Fig. 4D**), **MuSK** (0.47 ± 0.12-fold; **Fig. 4E**) and **Desmin** (0.36 ± 0.11-fold; **Fig. 4F**). Together, these findings demonstrate that silicone permeates exert direct myotoxic effects on skeletal muscle cells, characterized by reduced viability and disruption of the molecular program associated with myotube maturation, while preserving the initial fusion process.

**Figure 4.**
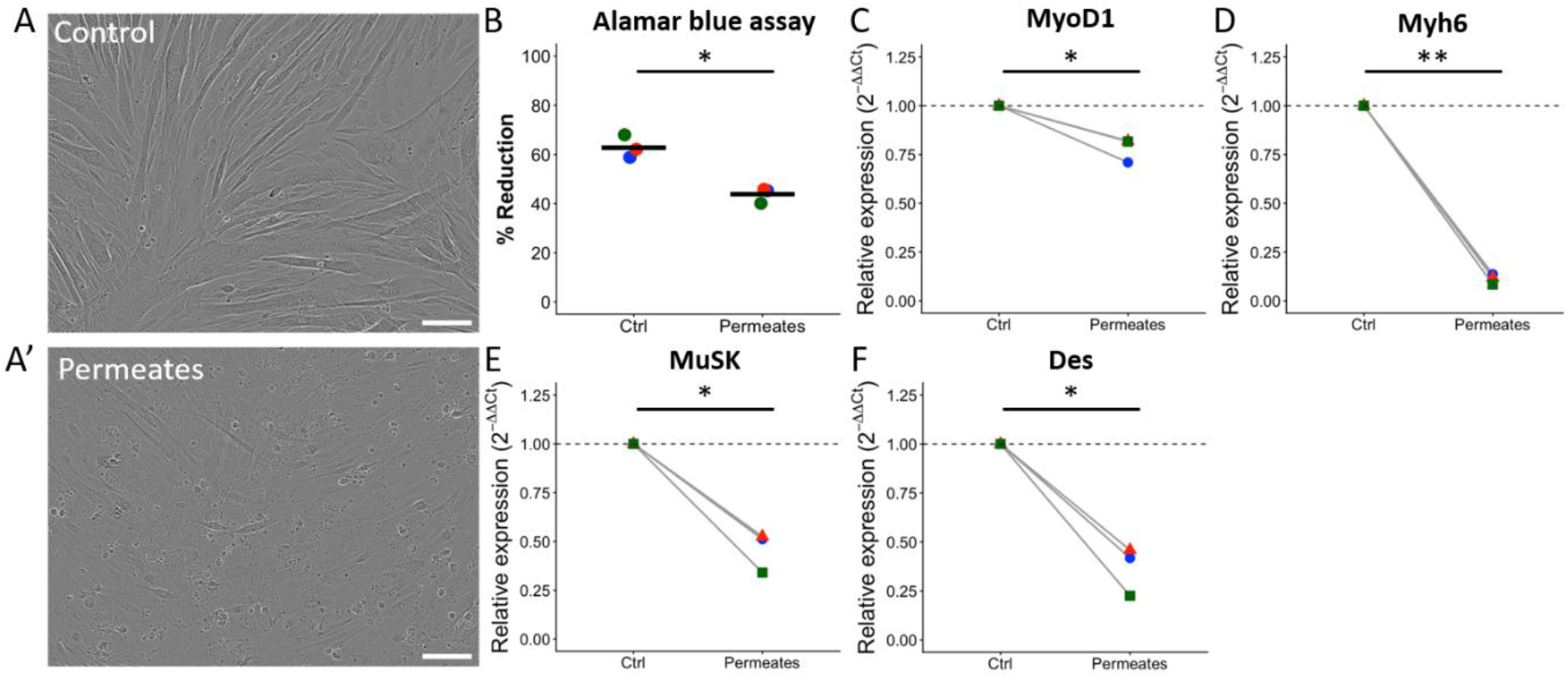
Silicone permeates impair C2C12 myotube viability and disrupt the myogenic transcriptional program. C2C12 muscle cells were differentiated for 6 days in control medium or in the presence of silicone permeates. (A) Representative phase-contrast images of C2C12. Scale bars, 100 µm. (B) Cell metabolic activity assessed by the Alamar Blue reduction assay. (C) RT-qPCR analyses of myogenic marker expression. The 3 independent biological replicates are shown in different colours. * p < 0.05; ** p < 0.01.

#### 2.3 Silicone exposure induces transcriptional remodeling of lipid metabolism and lipoprotein-associated pathways

A focused analysis of lipid metabolism-related genes identified 703 DEGs, including 344 upregulated and 359 downregulated (**Fig. 5A**). Enrichment analysis of GO biological processes within the lipid transport category revealed a broad spectrum of lipid-related functions, comprising lipoprotein transport (23.8%), cholesterol transport (18.8%), general lipid metabolism (15.0%), LDL/HDL receptor-mediated metabolism (12.5%), fatty acid metabolism (12.5%), regulation of lipid metabolism (10.0%), and specialized lipid transport and immune-related processes (7.5%) (**Fig. 5B**).

**Figure. 5.**
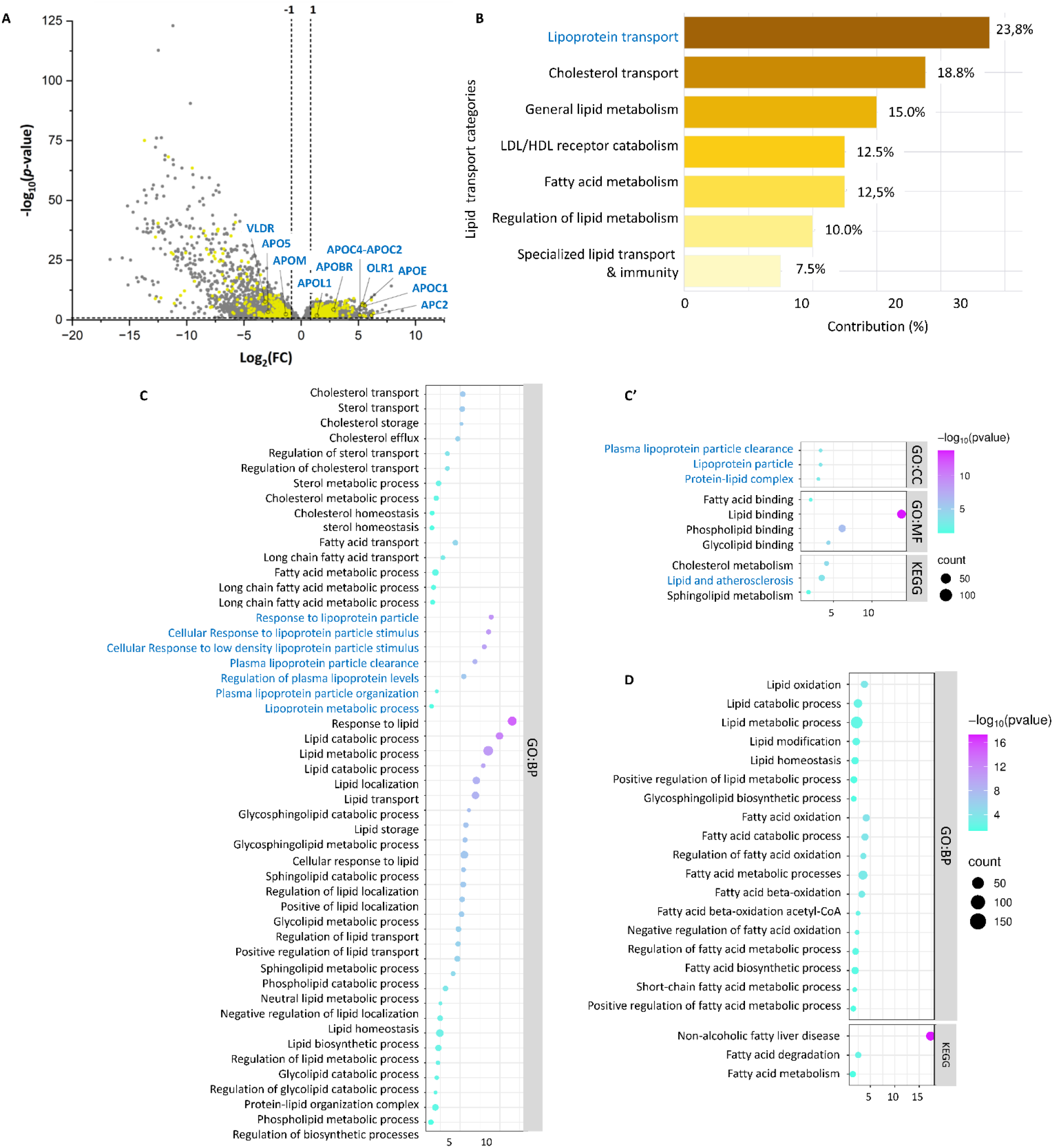
Silicone exposure following breast implant rupture dysregulates lipid metabolism and lipoprotein-related pathways in periprosthetic tissues. (**A**) Volcano plot showing differentially expressed lipid-related genes, with lipoprotein-encoding genes highlighted. (**B**) Relative contribution of lipid-related biological processes, expressed as the percentage of differentially expressed genes assigned to each functional category. (**C-D**) Bubble plot of significantly enriched lipid-related pathways, highlighting lipoprotein-mediated biological processes, with (**C-C’**) and (**D**) corresponding to the pool of up- and downregulated genes. Bubble color represents statistical significance (−log₁₀ P value), and bubble size corresponds to the number of associated genes.

Analysis of GO terms revealed enrichment of lipid regulatory processes, including cholesterol, sterol and (glyco)sphingolipids homeostasis, lipid transport, localization, and responses to lipid-derived signals (**Fig. 5C**). Remarkably, lipoprotein-related functions were prominently represented, with enrichment of GO terms associated with cellular response to lipoprotein, plasma lipoprotein clearance, regulation of plasma lipoprotein levels, lipoprotein organization, and lipoprotein metabolic processes. In parallel, GO cellular component analysis highlighted increased representation of lipoprotein-associated structures, including plasma lipoprotein particles and protein-lipid complexes, suggesting remodeling of lipoprotein-related pathways and lipid transport processes following silicone exposure (**Fig. 5C and 5C′**). Consistent with these findings, KEGG pathway analysis identified *Lipid and atherosclerosis* as one of the most significantly enriched pathways, reflecting coordinated regulation of genes involved in lipoprotein metabolism, lipid transport, and lipid homeostasis (**Fig. 5C′**). Examination of individual differentially expressed genes (DEGs) further supported extensive remodeling of lipid and lipoprotein-associated pathways. Among the downregulated genes, *VLDLR*, *APOL5*, and *APOM* were significantly decreased, along with multiple genes involved in fatty acid metabolism, indicating altered fatty acid handling (**Fig. 5D**). Conversely, several apolipoprotein-related genes, including *APOL1*, *APOC1*, *APOC2*, *APOE*, *APOBR*, *OLR1*, and the *APOC4–APOC2* locus, were markedly upregulated (**Fig. 5A**), suggesting activation of lipoprotein-associated transcriptional programs. The coordinated enrichment of lipoprotein-related pathways and altered expression of apolipoprotein genes suggest a shift toward enhanced lipoprotein recognition and transport programs, accompanied by altered lipid metabolic and catabolic processes. Finally, beyond lipoprotein metabolism, the enrichment of sterol and cholesterol together with (glyco)sphingolipid-associated pathways points toward a broader signaling network. Given the central role of cholesterol trafficking in lipoprotein metabolism and macrophage foam cell formation (observed after silicone exposure, **Fig. 3Ab**), these alterations may reflect changes in cellular lipid handling and responses to the accumulation of hydrophobic material. In parallel, alterations in (glycol)sphingolipid-associated pathways suggest potential effects on membrane dynamics and inflammatory signaling. Together, these findings highlight a complex transcriptional reprogramming of lipid-associated functions following silicone exposure, characterized by enrichment of lipoprotein-related pathways and altered expression of genes involved in cholesterol, fatty acid and (glyco)sphingolipid metabolism.

## Discussion

The migration of LMWS through intact silicone elastomer shells, commonly referred to as “silicone gel bleed” in breast implants, has been recognized for several decades[24–27]. However, the molecular composition of released silicone species under physiologically relevant conditions remains poorly defined. To address this knowledge gap, we analyzed an aqueous extract containing the readily releasable fraction of LMWS generated from commercially available silicone implant gel and characterized this mixture using the complementary analytical strengths of gel permeation chromatography (GPC) and gas chromatography–mass spectrometry (GC-MS). GPC provides information on apparent molecular size distribution and enables comparison with reference materials but cannot provide unequivocal structural identification. GC-MS complements this approach by chromatographically separating siloxanes and identifying them via mass spectral matching. Together, these methods provide robust characterization of released silicone species from our *in vitro* gel bleed model solution. Our analysis detected D4, D5 and D6, consistent with cyclosiloxanes reports in human blood[28] and peri-implant tissue[29,30]. Although our experimental system differs from the clinical environment, the detection of these compounds is therefore consistent with previous observations in human samples. Their detection in the present *in vitro* model is therefore consistent with observations made in clinical samples. Critically, we also identified L2, D3, D7, D8, and a putative D9, compounds rarely examined in biological studies. This broader molecular weight distribution (L2–D9) challenges the conventional focus on D3–D6, as many targeted methods prioritize lighter, more volatile siloxanes due to increasing analytical sensitivity constraints as molecular weight increases. The tentative detection of D9 suggests that the released silicone fraction may comprise a more chemically diverse oligomeric mixture than is generally appreciated. Furthermore, the lower volatility and increased hydrophobicity of D7–D9 may influence tissue retention, transport, and cellular uptake, particularly given reports of silicone in distant tissues post-implantation. Although these physicochemical properties provide a plausible basis for altered biological behavior, direct experimental evidence regarding the biodistribution of these higher-molecular-weight species remains limited. Low D3-D5 levels in extraction controls were expected, as cyclic siloxanes are known laboratory contaminants [20–22]. However, the elevated signals in gel-infused medium confirm an implant-derived source, warranting further characterization of the gel bleed signature. Our GC-MS analysis was semi-quantitative, designed for identification rather than absolute quantification. Future studies using isotope-labeled internal standards will enable accurate determination of release kinetics and concentrations. Nevertheless, our data demonstrate that commercial silicone gels release multiple cyclic siloxanes (L2-D8, plus putative D9) under physiological conditions, supporting the view that gel bleed involves a heterogeneous mixture rather than only light volatile siloxanes. Finally, it should be emphasized that the present study characterizes the detectable silicone-derived species released under the applied extraction protocol rather than the complete chemical composition of clinical gel bleed.

Previous studies have more focused on individual low-molecular-weight siloxanes (LMWS), often using experimentally high concentrations that are unlikely to reflect clinically relevant exposure levels, and have generally been limited to the cyclic species D4–D6, based on the assumption that their smaller molecular size makes them the predominant siloxanes released from silicone implants. Although individual compounds can exhibit variable cytotoxicity and induce apoptosis [31], this single-focused approach overlooks the cumulative biological effects of the complex siloxane mixtures released from silicone implants. Differences in molecular size, hydrophobicity, volatility, and likely cellular uptake among individual siloxanes may influence their partitioning, bioavailability, and cellular interactions, resulting in biological responses that extend beyond those of any single compound. To our knowledge, only one *in vivo* study has evaluated such a representative siloxane blend, demonstrating in *C. elegans* that gel bleed exposure reduces brood size and impairs progeny mobility[19]. Together, these findings emphasize that while the biological effects of individual LMWS remain important to characterize, understanding the cumulative effects of complex siloxane mixtures is essential for accurately assessing the biological impact of implant-derived silicone exposure.

Musculoskeletal manifestations, including myalgia, weakness and increased fatigability, are frequently reported by women with silicone breast implants[32–34]. Although these symptoms have mainly been discussed in the context of chronic inflammation, our findings provide convergent histological, transcriptomic and *in vitro* evidence that implant-derived LMWS directly affect skeletal muscle and the neuromuscular unit. Histological analysis revealed skeletal muscle fibers intimately embedded within the periprosthetic capsule, creating a close interface between silicone-exposed tissue and the adjacent pectoral muscle. This unexpected anatomical integration likely explains the prominent muscle-related signature detected in the periprosthetic RNA-seq dataset. Silicone-exposed tissues displayed coordinated downregulation of myogenic regulators, including *MYOD1, MYOG, MYF5, MYF6, MEF2C* and *MEF2D*, together with structural and contractile genes encoding desmin, myosins, troponins, tropomyosins and sarcomeric proteins. Moreover, our *in vitro* experiments support a direct contribution of LMWS to these alterations. Despite relatively preserved initial myoblast fusion, exposure to a silicone implant-derived L2-D9 permeate blend reduced cellular metabolic activity and induced progressive myotube degeneration. Consistent with the human transcriptomic dataset, LMWS exposure downregulated *MyoD1, Myh6, Des*, and *MuSK*, indicating that C2C12 cells recapitulate key molecular alterations observed in human tissues, and, therefore, represent a relevant model for investigating the direct effects of silicone-derived LMWS on skeletal muscle. The reduced expression of myogenic markers observed *in vivo* may reflect direct muscle-cell toxicity and impaired survival. Importantly, the absence of immune cells and exogenous cytokines in the *in vitro* model indicates that the observed effects result from a direct action of LMWS on muscle cells rather than from secondary inflammatory mechanisms, although inflammation may further potentiate these effects *in vivo*. The transcriptional alterations also extended to excitation–contraction coupling and neuromuscular-junction pathways. Dysregulated genes included *CACNA1S, RYR1, ATP2A1, CASQ1, STAC3, SCN4A* and *BIN1*, which regulate membrane excitability, calcium release and reuptake, and coupling between transverse tubules and the sarcoplasmic reticulum. Their coordinated alteration may impair calcium homeostasis and contractile efficiency, providing a plausible molecular basis for weakness, fatigability and cramps described by women with silicone breast implant. In parallel, genes involved in postsynaptic specialization and NMJ maintenance were affected, including *MUSK, LRP4, DOK7, RAPSN, ACHE, COLQ, CHRND* and *CHRNB1*. The agrin–LRP4–MuSK–DOK7 pathway is essential for acetylcholine-receptor clustering and stabilization of the postsynaptic membrane[35]. The downregulation of *MuSK* in both human tissues and C2C12 monocultures is therefore particularly relevant, as it indicates that LMWS can directly alter the muscle-side molecular program required for NMJ organization. This finding further supports the hypothesis that LMWS exposure affects the neuromuscular system. (**Fig. 6**). These results collectively suggest that the postsynaptic compartment is a potential target of silicone exposure. Finally, the dysregulation of genes associated with motor axons, Schwann cells and myelin suggests that the effects may extend beyond muscle fibers. Peripheral myelin is produced by Schwann cells and is essential for electrical insulation, trophic support and rapid motor-axon conduction[36]. Its exceptionally high lipid content makes it potentially vulnerable to lipophilic xenobiotics. In our dataset, multiple genes associated with Schwann cell biology and myelin maintenance (*GAL3ST1, GJC3, LAMA2, SCHIP1*) and mitochondrial quality control (*PRKN, PINK1, OPTN*) were dysregulated. Notably, the dysregulation of *PINK1*, *PRKN*, and *OPTN* suggests impaired mitochondrial homeostasis, a critical process for motor neuron maintenance and a hallmark of neurodegenerative disorders, including amyotrophic lateral sclerosis (ALS). These results support the hypothesis that LMWS may disturb peripheral glial or myelin homeostasis. Similar effects on lipid-rich neural structures have been reported for other persistent xenobiotics, including PFAS and microplastics[37]. Together, these findings support an integrated model in which LMWS may simultaneously impair myogenic maturation, excitation–contraction coupling and postsynaptic NMJ organization, with a possible additional effect on Schwann cells and peripheral myelin. These mechanisms may act synergistically, as direct myofiber toxicity could increase vulnerability to altered innervation, while impaired neuromuscular signaling could promote functional denervation and muscle loss. This model provides a plausible framework for implant-associated fatigue, weakness, myalgia and cramps, while dedicated functional studies will be required to confirm NMJ dysfunction and peripheral myelin injury.

**Figure 6.**
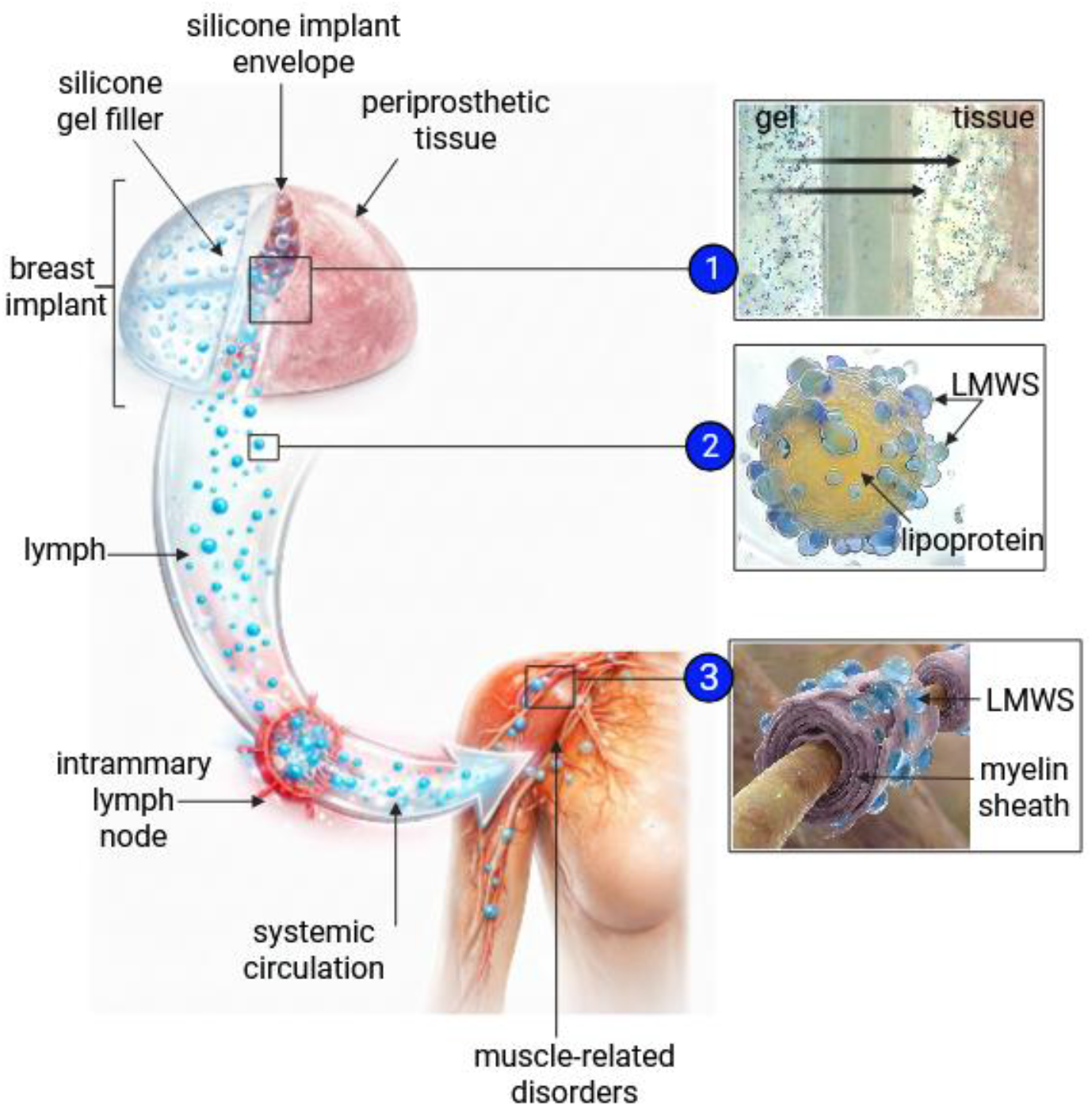
Hypothetical model of systemic LMWS silicone transport. (**1**) **Local exposure**: Periprosthetic tissues are exposed to low-molecular-weight silicones (LMWS) due to chronic silicone bleed from implants. (**2**) **Lymphatic transport**: LMWS associate with lipoproteins and are systematically transported via lymphatic circulation, occasionally accumulating in regional mammary lymph nodes. (**3**) **Systemic distribution**: In blood circulation, LMWS-lipoprotein complexes enable distribution to distant lipophilic organs, such as muscle, where hydrophobic LMWS may associate with myelin-rich lipid structures, potentially altering motor-neuron axis function.

Implant-derived silicone dissemination is supported by the detection of silicone in regional lymph nodes [38,39] and other distant lipophilic organ including liver, spleen or brain [5,7]. However, the pathways and extent of this dissemination remain controversial[40,41]. Due to their hydrophobic properties, silicone-derived species are likely to associate preferentially with lipid-rich environments rather than remain freely solubilized in aqueous fluids, suggesting another alternative supporting silicone transport. Accordingly, lipid-associated carriers, including lipoproteins, emerge as plausible mediators of silicone redistribution and systemic dissemination. Our RNA-seq data provide molecular support for this hypothesis by identifying coordinated transcriptional changes in pathways involved in lipid handling, including lipid metabolism, transport, and homeostasis, particularly those related to lipoproteins and cholesterol, in silicone-exposed periprosthetic tissues. Histologically, tissues adjacent to ruptured implants exhibited a palisade-like structure of lipid-laden macrophages (**Fig. 3Ab**), consistent with the engagement of lipid-associated inflammatory processes following silicone exposure. In parallel, the altered expression of sphingolipid- and glycosphingolipid-associated pathways may contribute to an immunometabolic phenotype resembling that observed in lipid-loaded macrophages, in which membrane lipid composition and innate immune signaling are closely interconnected[42–44]. This foam cell-like macrophage phenotype parallels processes observed in atherosclerotic plaque development, where macrophages internalize modified lipoproteins through scavenger receptors and Toll-like receptor pathways[45]. Consistent with this parallel, RNA-seq analysis revealed enrichment of lipoprotein recognition, scavenger receptor, and Toll-like receptor signaling pathways (described in [4]), together with altered gene expression related to lipid metabolism and transport. These findings support the hypothesis that silicone dissemination may partially exploit endogenous lipid-handling mechanisms, whereby macrophage lipid-handling pathways may contribute to the cellular uptake and sequestration of silicone-associated material and potentially participate in its redistribution. This interpretation is further supported by previous reports describing intracellular lipid accumulation and foam cell-like phenotypes in silicone-exposed tissues[46], as well as lipid infiltration associated with implant degradation and visible yellow discoloration of explanted devices[47–50]. Moreover, altered oxylipin profiles have recently been reported in patients with breast implant-associated systemic symptoms, further suggesting that dysregulated lipid mediator pathways may contribute to the biological response to silicone exposure[50]. Lipoproteins are well acknowledged for their involvement in the systemic transport and body distribution of hydrophobic molecules, such as postprandial lipids[51]. In line with this hypothesis, analogous pathways have been documented for other hydrophobic xenobiotics, such as volatile silicones in cosmetics [52] and persistent contaminants like PFAS [53] which associate with plasma lipoproteins and promote foam cell formation. This convergence with physiological transport mechanisms could explain how local silicone release reaches distant lipophilic structures, such as the neuromuscular junction, providing a mechanistic framework for silicone-associated myogenic alterations (**Fig. 6**). However, although our findings support a potential role of lipid-handling pathways in silicone redistribution, formally establishing systemic silicone dissemination will require addressing persistent limitations in tissue accessibility, standardized models, and silicone detection methods, which, to date, have hindered definitive conclusions[54].

## Conclusions

In conclusion, RNA sequencing of periprosthetic tissues associated with silicone implant rupture revealed coordinated alterations in myogenic and neuromuscular signaling, together with dysregulation of lipid metabolism and lipoprotein-associated pathways. These findings were supported by histological evidence of close interactions between periprosthetic tissue and skeletal muscle and by chemical characterization of the low-molecular-weight fraction released from implant gel through “gel bleed”. *In vitro*, exposure of differentiating muscle cells to the implant-derived fraction reduced cell viability by up to 30% and altered selected myogenic and neuromuscular genes identified in the clinical analysis, even in the absence of exogenous inflammatory stimulation. Together, the convergence of clinical, histological, chemical, and experimental findings supports the biological relevance of silicone gel bleed beyond a material-integrity phenomenon and provides evidence that its low-molecular-weight constituents can exert biological effects on skeletal muscle cells.

The concomitant alterations in lipid metabolism and lipoprotein-associated pathways provide a biologically plausible framework for investigating the cellular handling of hydrophobic silicone-derived species and their potential tissue distribution. However, our data do not establish a direct physical association between circulating siloxanes and lipoproteins, nor do they demonstrate systemic transport to distant tissues. Further studies will therefore be required to determine whether lipid-handling mechanisms contribute to the distribution and persistence of low-molecular-weight silicone species in vivo. Despite the limited clinical cohort (n = 11) and the preliminary nature of the mechanistic investigation, this materials-to-biology approach provides a basis for further investigating how silicone gel bleed may contribute to tissue responses following implant exposure, including musculoskeletal manifestations.

## Acknowledgements

We gratefully acknowledge the Agence Nationale de la Recherche, grant “Safe-implant” - ANR-22-CPJ1-0077-01), the Centre National de la Recherche Scientifique, and the Ministère de l’Enseignement Supérieur et de la Recherche for their financial support. We thank Cécile Joyeux of the molecular analysis platform at LIMA for the GC-MS analyses, BioRender (https://biorender.com) for the illustration software, Illustrae (https://illustrae.co/) for LMWS structure illustrations, and Adobe Firefly (https://www.adobe.com/firefly) for AI-assisted image generation.

## Author Contributions

I.B.: Conceptualization, supervision, writing-original draft, literature review, data curation, figure preparation, funding acquisition, RNA sequencing development and analyses, generation of original data.

N.C. and E.R: figure preparation, data analysis (RNA sequencing on muscle and *in vitro* experiments), literature review, and writing, review and editing draft.

E.R., H.M., C.M., and A.P.: Conceptualization, figure preparation, and data analysis (GPC and GC–MS).

I.P., F.M and F.B: Clinical sample collection and tissue collection storage in biobank

All authors: Contributed to the conceptual development of the study, the synthesis and interpretation of the literature, and the critical revision of the manuscript. All authors approved the final version of the manuscript and agree to be accountable for all aspects of the work.

## Data availability

The RNA-seq data generated in this study are not publicly available because they form part of an ongoing study. The data supporting the findings of this study are available from the corresponding author upon reasonable request.

## Ethics and integrity statement

The authors declare no competing financial or non-financial interests. Funding sources supporting the authors’ research activities are acknowledged separately where applicable. Permissions for reproduction of previously published figures or materials have been obtained from the respective copyright holders where required. This article reports findings from a clinical trial that has been partially published previously (DOI: 10.1016/j.biomaterials.2024.123025). The study was registered on ClinicalTrials.gov (Identifier: NCT06414785) and was conducted in accordance with the approval of the local Institutional Review Board/Ethics Committee. Written informed consent was obtained from the patient prior to participation.

